# Tropical Forest Soil Microbiome Modulates Leaf Heat Tolerance More Strongly Under Warming than Ambient Conditions

**DOI:** 10.1101/2024.12.09.626610

**Authors:** Gabriela Hernandes Villani, Iana F. Grullón-Penkova, Parker Bartz, Joel Masanga, Jesse R. Lasky, Sasha C. Reed, Molly A. Cavaleri, Tana E. Wood, Benedicte Bachelot

## Abstract

Tropical forests are increasingly threatened by climate change. Yet, it is still unclear how tropical plants respond to increasing temperatures. Leaf heat tolerance (LHT) in tropical plants is often at its upper limit, suggesting that climate change might negatively impact tropical forests. We hypothesized that intraspecific variation in this leaf trait might be associated with changes in the soil microbiome, which might also respond to climate. Specifically, we hypothesized that warming would increase LHT through changes in the soil microbiome: this study combined an *in-situ* tropical warming experiment with a shade house experiment in Puerto Rico. The shade house experiment consisted of growing seedlings of *Guarea guidonia*, a dominant forest species, under different soil microbiome treatments (reduced arbuscular mycorrhizal fungi, reduced plant pathogens, reduced microbes, and unaltered) and soil inoculum from the field experiment. Heat tolerance was determined using chlorophyll fluorescence (*F_V_/F_m_*) on individual seedlings in the field and on pooled seedlings by pot in the shade house. We sequenced soil fungal DNA to analyze the impacts of the field and shade house treatments on the soil microbiome.

In the field, seedlings from ambient temperature plots showed higher *F_V_/F_m_* values under high temperatures (0.648 at 46 °C and 0.067 at 52 °C) than seedlings from the warming plots (0.535 at 46 °C and 0.031 at 52 °C). In the shade house, the soil microbiome treatments, which significantly influenced the fungal community composition, significantly influenced LHT. Reduction in fungal pathogen abundance and diversity altered *F_V_/F_m_* before *T50* for seedlings grown with soil inoculum from the warming plots but after *T50* for seedlings grown with soil inoculum from the ambient plots.

Our findings emphasize that the soil microbiome might play an important role in modulating the impacts of climate change on plants. Understanding and harnessing this relationship might be vital for mitigating the effects of warming on tropical forests, emphasizing the need for further research on microbial responses to climate change.

## Introduction

Tropical forests are biodiversity hotspots, hosting approximately 70% of all species on the planet but covering just 15% of land surface (Pan et al., 2013). In addition, this biome acts as a carbon sink, absorbing and storing vast amounts of atmospheric carbon dioxide through photosynthesis, thus mitigating the impacts of climate change (Le Quéré et al., 2018). However, climate change threatens this crucial role, and tropical soils may be shifting to a carbon source in the atmosphere (Nottingham et al., 2019). During the last century, the global temperature has increased by 1 °C, and climate models predict that it will increase by 3.5–4.0 °C by the end of the century (Parry et al., 2007), which is a possible turning point in plant metabolic function (Doughty et al., 2023). However, little is known about how the most diverse biome on Earth responds to warming (Wood et al. 2019).

Leaf temperature is altered by sunlight, wind, and water availability and usually exceeds air temperature during periods of intense sunlight (Matsui & Eguchi, 1971; Perera & Eckard, 2019). Tropical plants tend to be at their maximal temperature tolerance (Mau et al., 2018; Tiwari et al. 2019; Sentinella et al., 2020), and their leaf temperature can be higher than air temperature by up to 10°C (Miller et al., 2021; Crous et al., 2023). Moreover, the acclimation capacity of many tropical plants is inadequate to maintain stable leaf thermal safety margins at higher growth temperatures, making them more susceptible to increased thermal stress (Tarvainen et al., 2022; Kullberg et al., 2024).

Leaf heat tolerance (LHT) is a functional trait describing the ability of a leaf to withstand high temperatures. This trait is determined by maximum quantum efficiency of the protein-complex Photosystem II (*F_V_/F_m_,* Baker and Rosenqvist, 2004; Krause et al., 2013), which is the ratio of maximum variable to maximum total fluorescence (Krause et al., 2010). As the leaf experiences high temperatures, *F_V_/F_m_* declines. Leaf heat tolerance is often defined by the parameters best describing this decline in maximum quantum efficiency of protein-complex Photosystem II, *T50,* and *Tcrit. T50* is the temperature where the efficiency of the protein-complex Photosystem II is reduced by 50% whereas *Tcrit* is the critical temperature at which leaves start to exhibit damage (Perez and Feeley 2020; Slot et al. 2021). High LHT is associated with high *T50* and *Tcrit*. In contrast, a low LHT corresponds to a low *T50* and *Tcrit*. Understanding variations in *F_V_/F_m_* and in these two commonly used LHT parameters may help predict how tropical forests respond to a warming climate.

Leaf heat tolerance exhibits high intraspecific variation (Teskey et al., 2015) caused by abiotic and biotic factors. Abiotic factors, such as prior exposure to elevated temperatures, result in heat stress priming and memory, which can enhance plant resilience to subsequent episodes of thermal stress (Baürle, 2016; Friedrich et al., 2019). Priming is an example of acclimation (Filaček et al., 2022), by which the plants respond to prior elevated temperatures by activating heat shock transcription factors, boosting their response to future elevated temperatures (Baürle, 2016). Heat stress memory happens when exposure to prior elevated temperatures leads to changes in the chromatin structure, which could result in future heat tolerance for the plants and their offsprings (Baürle, 2016).

Besides abiotic factors, plant tolerance to heat stress can be mediated by biota such as beneficial fungi, which can improve plants’ water uptake (Hubbard et al. 2014), or other beneficial soil microbes, which can boost plants’ responses to heat stress (Shekhawat et al., 2022). Thus, associations between plants with beneficial bacteria or fungi may increase LHT. Plant response to stress includes modifications in the levels, types, and regulation of primary and secondary plant metabolites (Ramakrishna & Ravishankar, 2011). Amino acids and sugars are primary plant metabolites vital for plant vitality and growth, Alkaloids and flavonoids are secondary metabolites known to serve defensive purposes (Ramakrishna & Ravishankar, 2011). Due to the costs of abiotic stress response, plants developed a delicate equilibrium between allocating resources to stress management versus growth and reproduction (Abd El-Daim et al., 2019). Yet, plant performance under heat stress tends to increase if the plants engage in beneficial associations (Abd El-Daim, 2019; Waqas, 2015). Associations with beneficial symbionts help the plants allocate more resources towards both primary and secondary compounds, thereby supporting both growth and stress tolerance (Abd El-Daim et al., 2019). As a result, plants associated with beneficial bacteria and fungi might exhibit higher LHT, characterized by high *F_V_/F_m_, T50* and *Tcrit*. Among the beneficial fungi, arbuscular mycorrhizal fungi (AMF) are important in tropical forests, as they form a symbiotic relationship with more than 80% of tropical plant species (Tedersoo et al., 2018). Studies have highlighted the role of AMF in plant nutrient uptake, including calcium (Sardans et al., 2023) which is associated with high LHT (Jiang & Huang 2011). Therefore, AMF might be an important player in the soil microbiome with the potential to enhance LHT.

In contrast, fungal pathogens might be associated with low LHT, characterized by low *F_V_/F_m_, T50* and *Tcrit*. Plant-pathogen interactions can be influenced by season, host plant, and environment (Elad & Pertot 2014; Velásquez et al. 2018), as well as the combined impact of these interactions (Nelson, 1994). An interaction with pathogens triggers defense pathways mediated by phytohormones, protein kinases, and programmed cell death in plants (Suzuki & Katano, 2018). As a result, plant’s ability to fight off pathogens may drain resources needed to maintain high levels of LHT. Alternatively, because of similarities in the signaling pathways involved in the response to pathogen attack and heat stress, one could expect higher LHT in plants interacting with pathogens than in healthy plants (Suzuki & Katano 2018).

The soil microbiome may also be directly affected by climate change (Cao et al., 2020; Nottingham et al., 2022). For example, in an experiment warming soil to +5°C AMF growth rates were increased for a few months, followed by a reduction in AMF diversity and a change in AMF community composition as the warming continued for three years (Cao et al., 2020). Similarly, bacterial diversity increased and peaked quickly before decreasing in a 2-year soil warming experiment in Panama (Nottingham et al., 2022). Finally, the life cycle of pathogens is influenced by temperature, which alters their survival and multiplication rates (Bale et al., 2002). High temperatures can boost or reduce the receptiveness of plants to a pathogen, depending on the host and the capacity of the pathogen to tolerate heat (Bale et al., 2002). Thus, climate change can alter the soil biota composition, which might influence plant LHT.

At the Tropical Responses to Altered Climate Experiment (TRACE) in Puerto Rico, early analyses of understory responses to warming highlighted interesting changes in biotic interactions due to altered climate (Bachelot et al., 2020). Seedling survival was increased in a +4 °C warming treatment when surrounded by high conspecific density without a decrease in aboveground natural enemies, suggesting potential long-term impacts on tropical wet forest diversity under future climate conditions (Bachelot et al. 2020; Alonso-Rodríguez et al. 2022).

In the present study, we sought to disentangle the effects of warming on competition, facilitation, mutualism, and antagonistic interactions among seedling demographics. We hypothesized that experimental warming regulates LHT by influencing the soil microbiome (Figure 1). Under this hypothesis, we predicted that; (i) LHT would vary between warmed and ambient plots due in part to inherent differences in microbial composition; and (ii) high relative abundances of fungal pathogens and beneficial communities like AMF would decrease and increase LTH, respectively. We tested this hypothesis using field and shade-house experiments.

**Figure 1.**
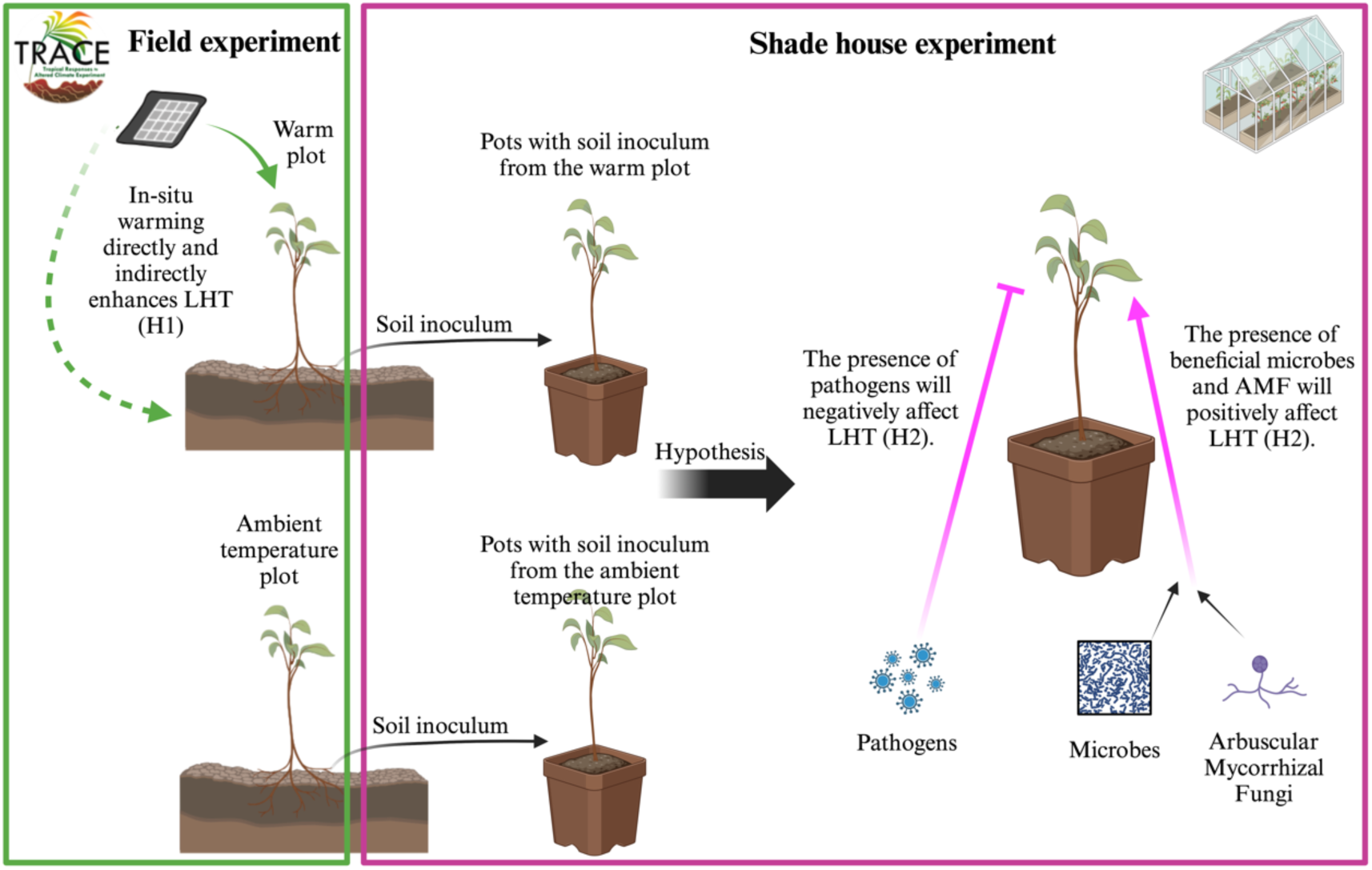
Graphical representation of hypotheses in the present study. Under favored hypotheses, experimental warming underlies variation in LHT by directly (plain line) impacting the plants and indirectly (dotted line) by regulating the microbiome under field (green: H1) and shade house (purple: H2) experimental conditions. Arrows represent predicted positive effects on LHT whereas perpendicular lines represent hypothesized negative effects on LHT.

## Methodology

### Site and TRACE field experiment

This study was conducted at the USDA Forest Service Sabana Field Research Station in Luquillo, northeastern Puerto Rico (18°18′N, 65°50′W). The area is described as a subtropical wet forest (Holdridge, 1967), typically aseasonal (Bouskill et al., 2016), and a secondary forest naturally recovered from pasture for almost 73 years (Kimball et al., 2018). The forest mainly comprises *Prestoea montana, Psychotria brachiata, Syzygium jambos,* and *Sloanea berteriana* (Cook et al., 2013). Rainfall is at least 200 mm per month and varies significantly throughout the year (Heartstill-Scalley et al., 2007), while the average annual temperature is 24 °C with low variation between months (4 °C, Kimball et al., 2018). The soil is described as Ultisol, with an elevated amount of weathered clay (Scatena, 1989).

The TRACE field experiment was established in 2015 to investigate understory plant and soil responses to +4 °C warming, using six 12 m² plots (Kimball et al., 2018). Three of these plots are heated using infrared heaters at 4 °C above the ambient level, suspended approximately 3.6 meters above the ground on crossbars. In contrast, the soil temperature sensors indicate that the soil in the warmed plots was about 1 °C higher than the temperature in the ambient temperature plots (Tunison et al., 2023). The remaining three serve as control plots, equipped with identical infrastructure but utilizing metal plates instead of heaters to simulate the physical presence of heating equipment without altering temperature. These plots are spaced about 10 meters apart in an open area without mature trees to accommodate the experimental setup (Bachelot et al., 2020; Alonso-Rodríguez et al., 2022; Kimball et al., 2018; Reed et al., 2020). The study’s warming treatment ceased on Sept 6, 2017, after 11.5 months, coinciding with Hurricane Irma approach 97 km north of Puerto Rico at Category 5 strength, followed by Hurricane María crossing the island on Sept 20, 2017, as a Category 4 storm (Yaffar et al.,2021). Warming was initiated again in September 2018 (Alonso-Rodríguez et al., 2022), being a 5-year warming experiment by the time of the data collection for this study which happened in July 2023.

### Field data collection

To assess the impact of increased temperatures on leaf heat tolerance (LHT), we focused on the dominant forest species, *Guarea guidonia*. This choice was informed by the species’ abundant seed supply and the availability of previous LHT data from earlier studies at the TRACE location (Carter & Cavaleri 2018; Carter et al., 2020). *Guarea guidonia*, commonly known as Muskwood, is an evergreen tree that can reach up to 23 meters in height, with a straight trunk up to 90 cm in diameter. *Guarea guidonia* is a late-successional broadleaved evergreen (Pennington et al., 2003).

In July 2023, we collected data from fully developed top leaves from two *Guarea guidonia* seedlings available per plot, which have been under heat treatment since their germination. We chose seedlings with similar heights (mean = 20.31 cm, range: 13 to 45 cm) and diameters (mean = 3.65 mm, range: 2.2 to 9.5 mm), and measured the height and diameter of each individual. We used a disk paper puncher (0.25 inches, Kawendite) to collect 18 leaf disks per plant and stored them in Ziploc bags for LHT assessment in the laboratory (see the LHT measurement section). Finally, with a sterilized metal spoon, we collected one soil sample from a soil core from each plot and kept them in sterile plastic bags in a −80 °C freezer until further analysis (see soil DNA analyses).

### Shade house experiment

We used a fully factorial design where 240 seedling pots were randomly assigned to one of four soil microbiome treatments with soil inoculum from either warmed or ambient plots. Pots were placed across four blocks, each assigned to low or high soil moisture treatment. For this study, we measured data on seedlings from the high soil moisture treatment to avoid confounding microbiome treatment and water stress effects of LHT. Each combination of treatment was replicated five times. Briefly *G. guidonia* seeds were collected in January 2023 from around the Sabana Field Research Station and their surface was sterilized using 5% sodium hypochlorite for 5 minutes as previously described (Sauer and Burroughs, 1986). The seeds were pre-germinated in sterile soil and transplanted (1, 3, and 5 seeds for low, medium, and high density, respectively) in pots with 250 g of oven-sterilized soil primed with 5 g of inoculum from the ambient or warmed plots (Figure 2). The four soil treatments were: unaltered microbiome, reduced AMF, reduced soil fungal pathogens, and reduced microbes. Reduced AMF was achieved by diluting 3.25 g of Banrot 40wp fungicide ((3-(2-methyl piperidine)-propyl - 3,4-dichloro benzoate) and 3.6 L of deionized (DI) water. Banrot is known for managing damping-off, root, and stem rot diseases instigated by *Pythium, Phytophthora, Rhizoctonia, Fusarium*, and *Thielaviopsis* (Prabhakaran & Dann, 2022). More importantly, Banrot is similar to Topsin-M which was found to significantly reduced AMF in the field (Wilson & Williamson, 2008). Subsequently, 60 mL of the diluted Banrot solution was poured into each pot and randomly assigned to the treatment. Reduced fungal pathogens were obtained by mixing 0.185 mL of Abound Flowable fungicide (Azoxystrobin: methyl(E)-2-{2-[6-(2-cyanophenoxy) pyrimidin-4-yloxy] phenyl}-3-methoxyacrylate) with 600 mL of DI water. Abound is a fungicide used for comprehensive disease management (Starkey et al., 2013). 2 mL of the diluted Abound solution was sprayed onto seedlings’ leaves in each pot assigned to this treatment. Finally, for the reduced microbe treatment, the soil inoculum was oven-dried for 48 h at 70 °C before pot filling. Both fungicide treatments (Banrot and Abound) were reapplied to respective treatments once every month.

**Figure 2.**
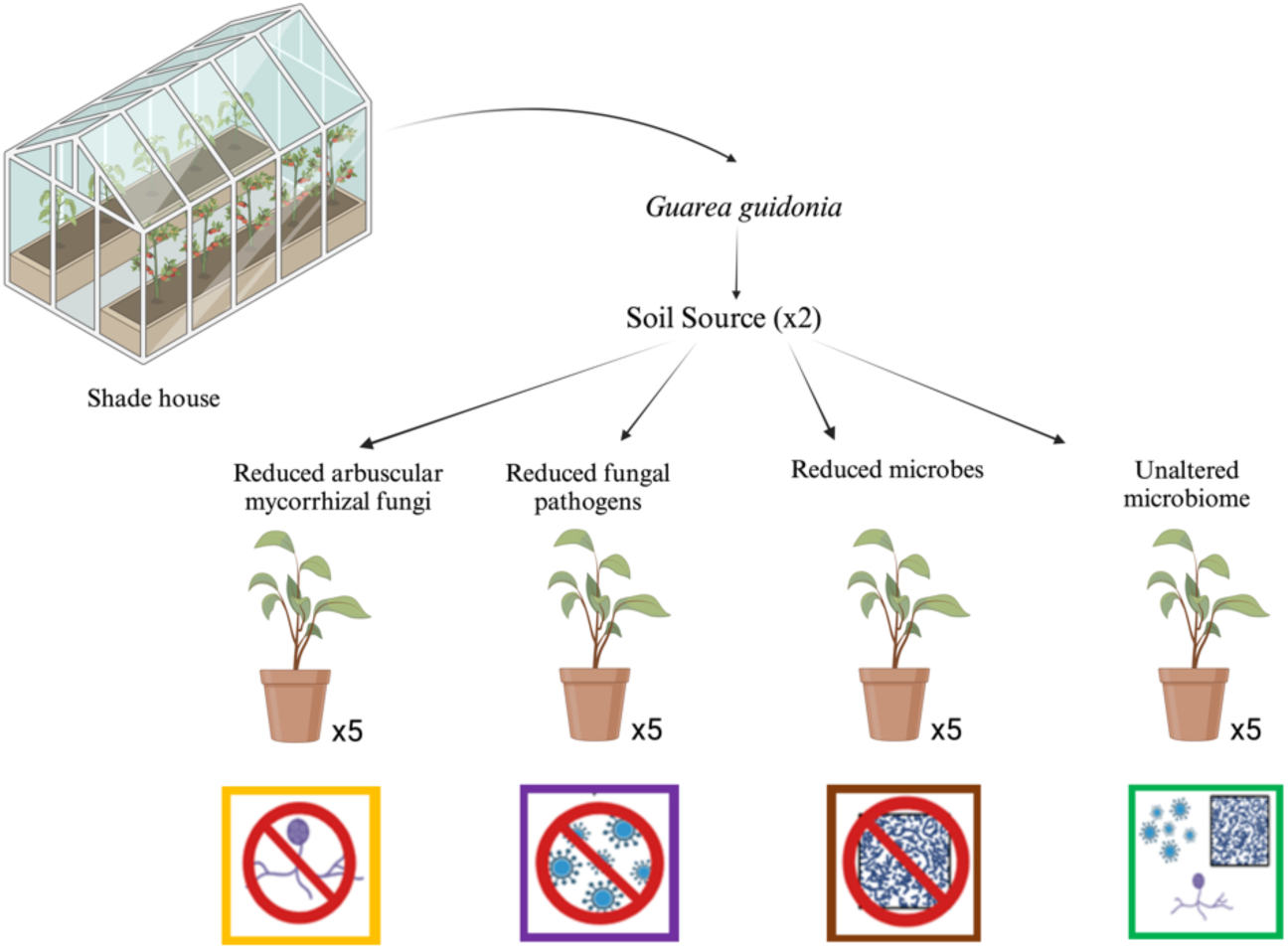
Schematic of the experiment. We grew *Guarea guidonia* in pots with soil inoculum from the warmed plot and the ambient temperature plot of five seedlings each. We exposed them to four different treatments (five replicates, reduced AMF (with microbes and fungal pathogens, in yellow), reduced fungal pathogens (with microbes and AMF, in purple), reduced microbes (with AMF and fungal pathogens, in brown), and unaltered microbiome, in green. Created with BioRender.com.

Seedlings in the shade house experienced low light conditions (shade was 60% and shade house was positioned in the shade of a building) and average temperature of 26 ± 3°C, which was similar to field conditions. Seedlings were allowed to grow for three months before LHT measurement. Three pots with the most leaf material available were selected from each of the five-pot replicates per treatment to collect leaf disks in July 2023. Each seedling from which leaf disks were collected was measured for height (mean = 11.55 cm, range: 5.1-18.5 cm) and diameter (mean = 3.32 mm, range: 1.9-3.8 mm). Using an ¼ inch disk paper puncher (Kawendite), 18 leaf disks per selected pot were collected, placed in a plastic bag, and immediately taken to the lab to measure LHT (see supplementary information Figure S1). We combined the 18 leaf disks from one pot into one measurement of LHT because individual seedlings did not have enough leaf material. Therefore, the shade house data is at the seedling pot level. Finally, we collected soil samples from each pot at the end of the shade house experiment and kept them in sterile bags in a −80 °C freezer until further analyses (see soil DNA analyses).

### Leaf heat tolerance measurement

Leaf disks were exposed to heat stress to measure LHT, following a method adapted from Krause et al. (2010). Leaf disks from both field and shade house experiments were collected early in the morning and immediately taken to the lab for preparation in Ziploc bags for LHT measurement. Approximately 30 minutes after collection, samples were ready to be incubated for 15 min in temperature-controlled water baths (water heater model ANOVA Sous vide 2.0, 3.0, precision +/- 0.2°C) set at one of five temperatures, target as common range for *T50* occurrence (Krauser *et al*, 2010, 2013): 46, 48, 50, 52, and 54° C; in addition, a sample was set at ambient temperature in the lab (25 °C) for control. Twenty-four hours later, leaf disks were placed in the dark for 15 min before *F_V_/F_m_* was measured using a fluorometer (model OS30p from Opt-Sciences). For each sample (individual plants in the field or pots in the shade house), *F_V_/F_m_* was measured using three leaf disks per temperature, totaling 18 leaf disks per sample.

*F_V_/F_m_* data were used to calculate several key parameters (*T50* and *Tcrit*) for each sample. *T50* is the temperature at which the potential quantum efficiency of Photosystem II (*F_V_/F_m_*) is reduced by 50% (Krause et al., 2010). *F_V_/F_m_* values after 24h of heat treatment is the parameter most closely associated with permanent leaf damage (dark coloration and necrosis, Krause et al., 2010). The parameter *b* represents the steepness of the decrease in *F_V_/F_m_* slope at *T50*. These parameters were obtained by fitting the *F_V_/F_m_* data to the following equation:

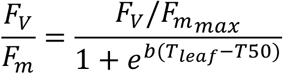

In this equation, *F_V_/F_m_*, _max_ is the upper horizontal asymptote representing *F_V_/F_m_* associated with healthy, non-stressed leaves. Finally, *T_leaf_* is the leaf disk’s incubation temperature (in °C). Once *T50* and *b* are estimated, the *Tcrit* parameter is obtained as the temperature where the line describing the slope of *F_V_/F_m_* declines at *T50* (*b*) intersects with the asymptotic line defined by *F_V_/F_m_*, _max_.

### Soil DNA extraction and sequencing

To characterize fungal community composition, we extracted and sequenced DNA from soil samples collected from warmed and control field plots as well as pots in the shade house. Briefly, a soil sample was scooped using a sterilized spoon at the center of each pot in the shade house and cores from the field, which were frozen at –80 °C in sterilized bags until, later, be subjected to DNA extraction using the PowerSoil Pro Kit (Qiagen, Germantown, MD, USA) according to the manufacturer’s instructions. Subsequently, the DNA extractions were sequenced via Illumina sequencing at the University of Minnesota Genomic Center (UMGC), using fungal-specific primers targeting the internal transcribed spacer 2 region (ITS4_Nextera and JL0015.8SR_Nextera). The University of Minnesota Genomic Center protocol includes a 3 min 95° C denaturation phase followed by a 20 s 98° C phase to activate the DNA polymerase, a 15 s 65.7 °C annealing phase, and a 45 s 72 °C elongation phase for 30 cycles, followed by a final 5 min 72 °C elongation (Gohl et al., 2016). To ensure data quality and correct sample differentiation, UMGC conducted preliminary quality control and demultiplexing. Sequencing data were analyzed using the Pete supercomputer at Oklahoma State University and the Mothur pipeline (Schloss PD et al., 2009). After detecting and removing chimeras, this pipeline generates operational taxonomical units (OTUs) by grouping sequences at 97% identity (Gweon et al., 2015) using the UNITE v9 database (Abarenkov et al., 2024). The resulting OTUs were taxonomically classified into phylotypes and putative guilds utilizing FUNGuild, a Python-based tool designed for ecological interpreting of fungal OTUs (Nguyen et al., 2016).

### Statistical analyses

To test whether warming influenced leaf heat tolerance in *G. guidonia* in the field (H1), we first tested how the raw data, the maximum quantum efficiency of PSII (*F_V_/F_m_*), differed between the ambient and warmed plots at each temperature (Slot et al., 2019), using a t-test. Then, we fitted mixed linear regressions to explain each estimated leaf heat tolerance parameter (*T50 and Tcrit*) as a function of warming treatment and seedling height as fixed effects and plots as a random effect.

To test whether the soil microbiome influenced leaf heat tolerance in the shade house (H2), we first investigated how *F_V_/F_m_* differed across biotic treatments at each temperature (Slot et al. 2019) using an analysis of variance (ANOVA) and Tukey post-hoc test to analyze which pairwise treatment differences were significant. This analysis was conducted within soil source. We then fitted a linear regression to explain each estimated leaf heat tolerance parameter (*T50 and Tcrit*) as a function of seedling average height, number of seedlings alive in the pot, microbiome treatment, soil source, and the interactions between soil source and microbiome treatment as fixed effects. Leaf heat tolerance parameters were obtained for a pot and not for an individual seedling (not enough leaf material) so we used the average seedling height as a covariate since leaf heat tolerance might change along ontogeny (Klockmann ad Günter, 2017). Finally, we used the uncertainty (standard error) from the fitted leaf heat tolerance parameters for each model as weights to account for errors in the response variable (heat tolerance parameters). Finally, we calculated estimated marginal means using the R package *emmeans* (lenth et al., 2022) for pairwise comparison across treatments and soil source in the shade house.

To test for differences in fungal composition, we used redundancy analyses to test if *in-situ* warming and soil microbiome treatments influenced the fungal community composition after verifying that fungal composition had homogeneous variances across plots and soil microbiome treatments. Furthermore, to identify how fungal classes differ among the treatments, we conducted an ANOVA followed by a Tukey post-hoc test to assess which fungal classes’ relative abundance differs across soil microbiome treatments.

All the analyses were conducted in R (version 4.2.1; R Core Team, 2022), using *nls.multsart* (Padfield & Matheson, 2018), *lmerTest* (Kuznetsova et al., 2017), and *vegan* (Oksanen et al., 2022) packages.

## Results

### Results from the Field

#### Warming lowers *F_V_/F_m_* in the field

As expected, *F_V_/F_m_* (Figure 3) was lower in seedlings from the warmed plots compared to ambient plots with mean values of *F_V_/F_m_* of 0.535 in the warmed plot and 0.648 in the ambient temperature plot for a 46 °C heat stress (t= 2.74, p=0.01) and 0.067 in the ambient temperature plot and 0.031 in the warmed plot for a 52 °C heat stress (t= 2.04, p=0.05). This result was reversed at ambient temperature (25°C), where *F_V_/F_m_* (Figure 3) was higher in seedlings from the warmed plots (t= - 2.69, p=0.011) with mean *F_V_/F_m_* values of 0.792 compared to seedlings from the ambient temperature plots with mean *F_V_/F_m_* values of 0.781.

**Figure 3.**
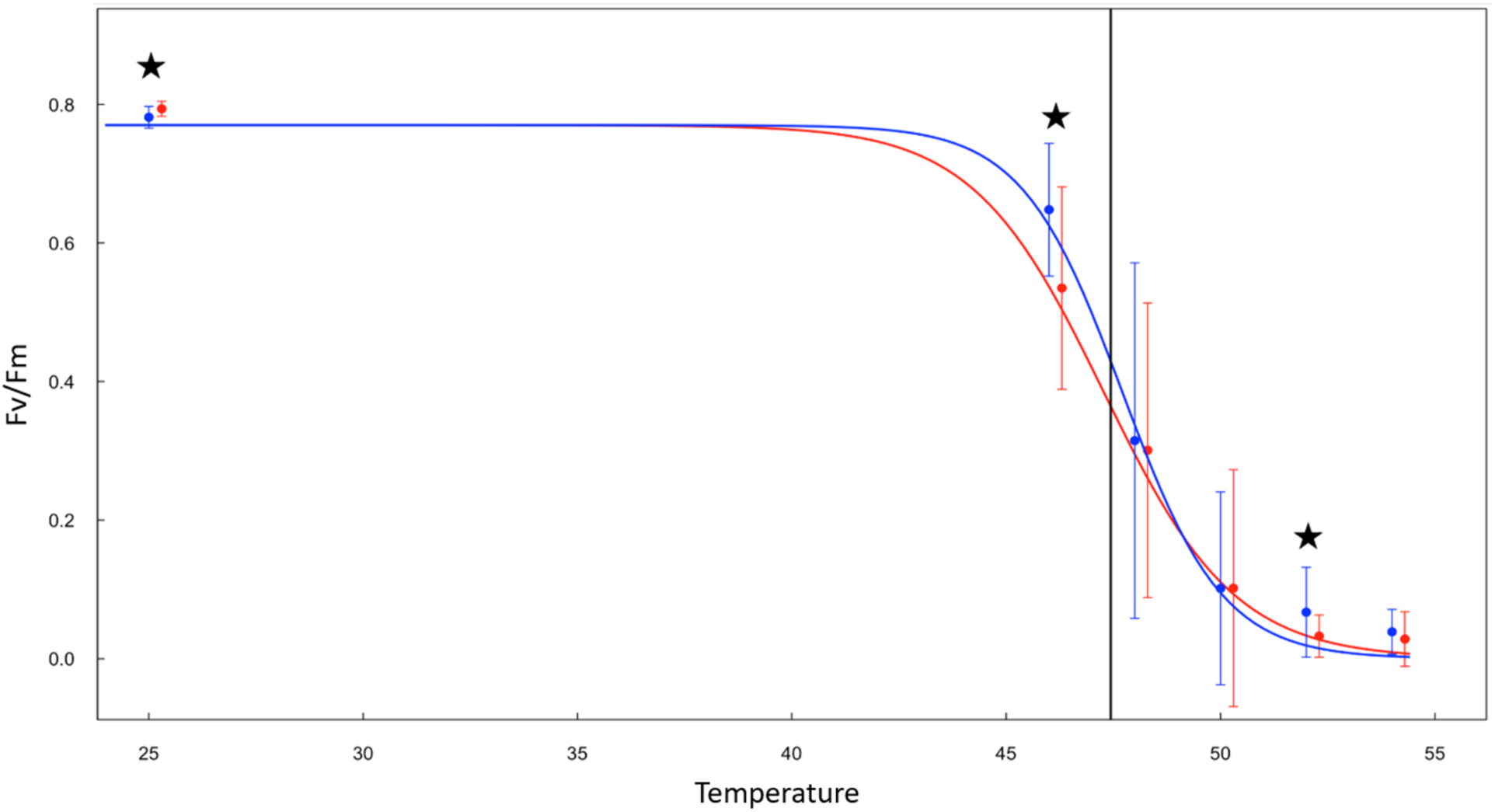
Mean *F_V_/F_m_* values according to the soil source from the plots (ambient (blue line) and warmed (red line)). The vertical bars show the *F_V_/F_m_* range at each temperature, and the dots show the mean values. The stars indicate significant differences (*p* ≤ 0.05) between the two soil sources at 25, 46, and 52°C according to the t-test. The black vertical line represents the mean T50 of seedlings in ambient and warmed plots.

Contrary to expectations, there was no significant difference in *T50* between seedlings grown in warmed versus ambient plots, nor were there significant differences in *Tcrit*.

#### Warming partially influences microbiome composition in the field

We hypothesized that the warming treatment might alter the microbial community structure. The analyses of the soil inoculum from the plots showed no significant effect of warming on overall fungal community composition, using the ANOVA test. The relative abundance of fungal classes most represented by pathogens such as *Dothideomycetes*, *Eurotiomycetes*, and *Sordariomycetes* was not significantly different between warmed and ambient plots. However, *Glomeromycetes*, which is an AMF class, is most present in the inoculum from the ambient plots, and *Archaeosporomycetes*, another AMF class, is just present in the inoculum from the warmed plot (Figure S2).

### Results from the shade house

#### Shade house treatment significantly altered the soil microbiome

While the fungal biomass, tested with the linear analysis, remained statistically consistent across treatments in both ambient and warmed plot soil sources (Figure S3), a significant difference in fungal DNA composition was identified across the microbiome treatments (p=0.001, F_3,82_=1.44, Figure 4A). Additionally, there was a significant interaction between soil source and microbiome treatments (p=0.009, F_3,82_=1.25). Further analyses across soil sources highlighted key fungal classes altered by the treatments, such as *Agariomycetes* (p=0.02, F_3,82_=3.67), *Sordariomycetes* (p=0.05, F_3,82_=2.69), and marginally significant for *Eurotiomycetes* (p=0.07, F_3,82_=2.39). However, the Tukey test only showed a significant difference, with values increasing respectively, for the *Agaricomycetes* class in the pairwise reduced AMF and reduced fungal pathogens (p=0.05, t=0.50), reduced AMF and reduced microbes (p=0.04, t=0.51), and marginally significant for unaltered and reduced AMF (p=0.07, t=15.21). Nevertheless, we observe a decrease in the log relative abundance of known fungal pathogenic classes such as *Dothideomycetes* in the treatments with reduced fungal pathogens and reduced arbuscular mycorrhizal fungi (AMF), and *Eurotiomycete* and *Sordariomycetes* in the treatments with reduced microbes, reduced fungal pathogens, and reduced AMF, compared to the unaltered treatment. Finally, *Glomeromycetes*, which contains AMF, is only present in the unaltered treatment (Figure 4B).

**Figure 4.**
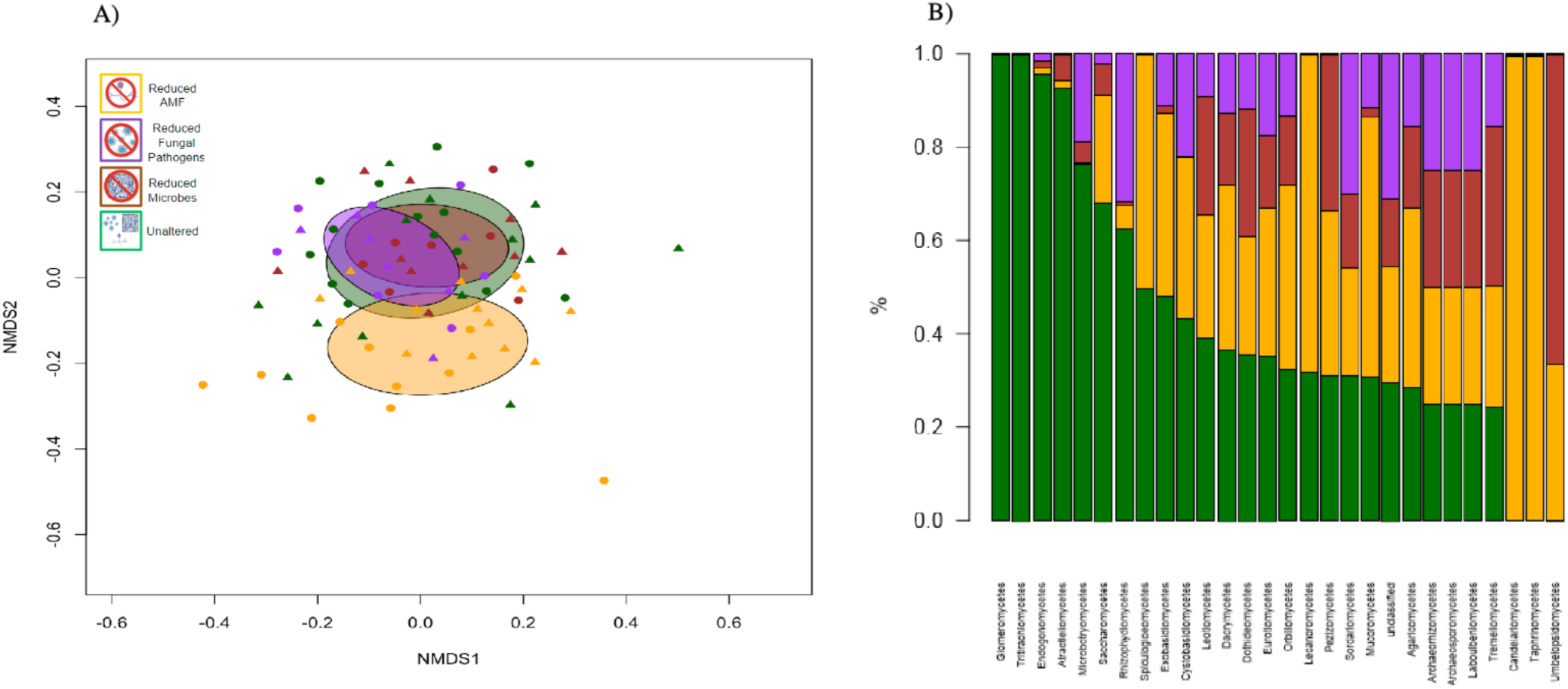
A) Non-metric multidimensional scaling to visualize the results of the redundancy analysis showing a significant effect of treatment (color of the symbols) and a treatment by soil source (circle: ambient plot; triangle: warmed plot) interaction on fungal composition. B) Log relative abundance (number of reads), standardized to 1, of fungal classes across the treatments in the shade house experiment, sorted from the highest to the lowest abundant in the Unaltered treatment. Reduced AMF (yellow), Reduced Fungal pathogens (purple), Reduced Microbes (brown), Unaltered (green).

#### The soil microbiome significantly influenced *F_V_/F_m_* and Tcrit in the shade-house experiment

*F_V_/F_m_* significantly differed across soil microbiome treatments. In the ambient plot soil source, soil microbiome treatment significantly influenced *F_V_/F_m_* for heat stress at 50 °C (p= 0.008, F_3,44_=0.04), 52 °C (p=0.031, F _3,44_=0.03), and 54 °C (p=0.072, F _3,44_=0.01) (Figure 5A). Plants grown with reduced AMF (p=0.02 t=-0.017) and reduced microbes (p=0.01 t=-0.028) had significantly higher *F_V_/F_m_* than plants grown with an unaltered microbiome for a heat stress at 50 °C. For heat stress of 52 °C (p = 0.02, t =0.26) and 54 °C (p=0.09, t =0.008), plants grown with reduced microbes showed higher *F_V_/F_m_* than those grown with reduced AMF.

**Figure 5.**
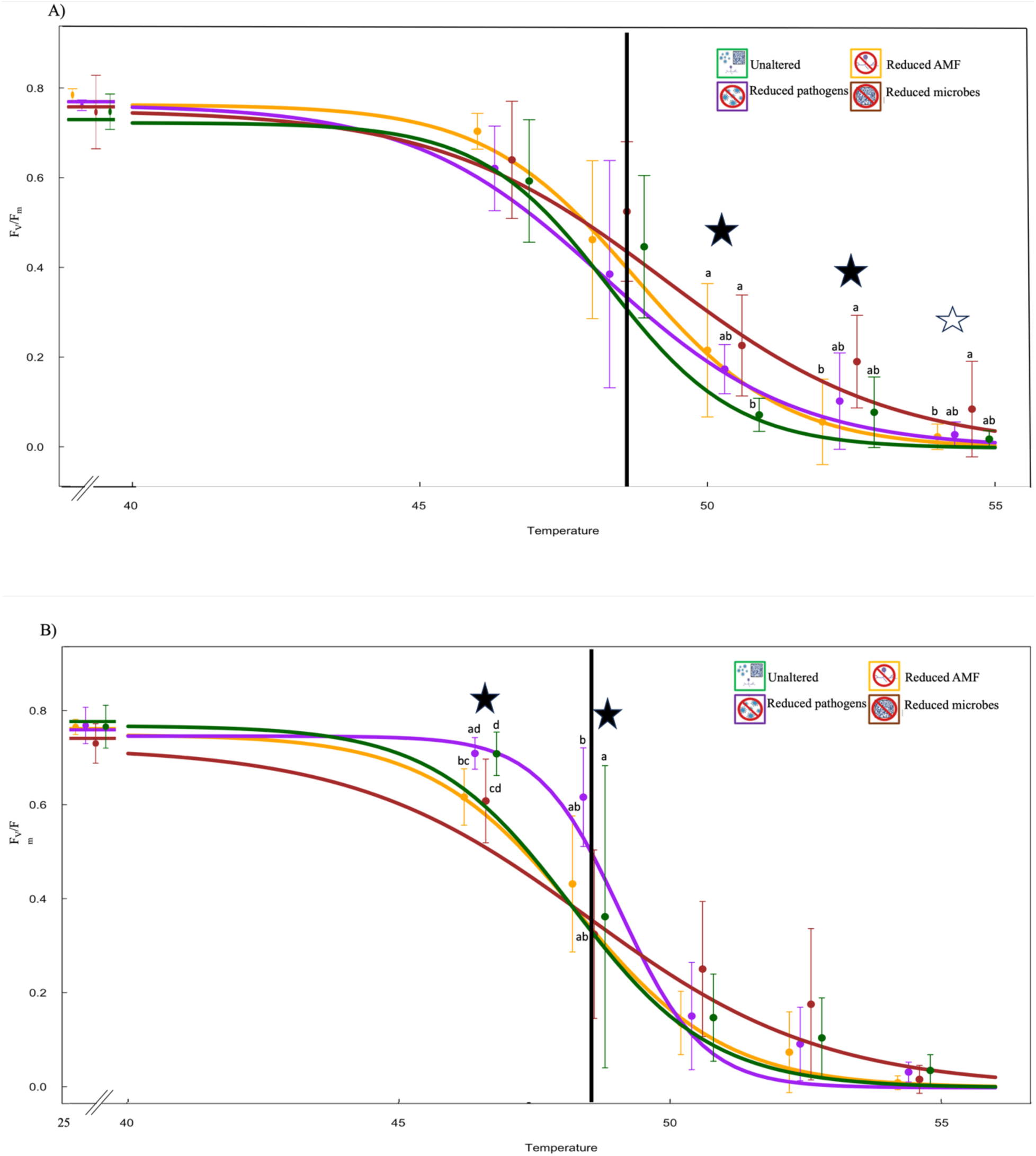
*F_V_/F_m_* mean values according to different treatments for ambient temperature (A) and warm (B) soil sources in the shade house. The filled stars indicate a significant difference (p-level, post-hoc test) among the treatments for heat stress at 50 and 52 °C for the ambient plot soil source and for heat stress at 46 and 48 °C for the warmed plot soil source. The empty star indicates a slightly significant difference (0.05≤p-value≤0.1) among the treatments for heat stress at 54 °C for the ambient temperature plot soil source. The orange line is T1(reduced AMF), the purple line is T2(reduced fungal pathogens), the brown line is T3(reduced microbes), and the dark green line is T4 (unaltered microbiome). The bars indicate the range. The black line indicates the mean *T50* value across treatments. The letters indicate which pairwise difference was statically significant.

In the warmed plot soil source, *F_V_/F_m_* significantly differed across soil microbiome treatments at 46 °C (p= 0.001, F _3,44_=0.02) and 48 °C (p=0.018, F _3,44_=0.14) (Figure 5B). The seedlings grown with reduced fungal AMF had lower *F_V_/F_m_* values than those grown with reduced pathogens (p=0.02, t=0.17) or with an unaltered microbiome (p=0.02, t=0.17) at 46 °C. Seedlings grown with reduced fungal pathogens had a significantly higher *F_V_/F_m_* value than seedlings grown with reduced microbes (p=0.01, t=-0.04) at both 46 °C and 48 °C.

Contrary to expectations, there were no significant differences in one of the leaf heat tolerance parameters (*T50*) among the different soil sources and treatments. Although, T50 was marginally higher in seedlings from the reduced pathogen treatment from the warmed plot soil source compared to seedlings from the reduced pathogen treatment from the ambient plot soil source (p=0.070, t= −1.96). The critical temperature at which heat stress initiates disruption of photosystem II (*Tcrit*) was significantly higher in the reduced fungal pathogen treatment (47.50 °C) from the warmed plot soil source than reduced pathogens in the ambient plot soil source (45.5 °C) (p=0.026, t= −2.48, Figure 6).

**Figure 6.**
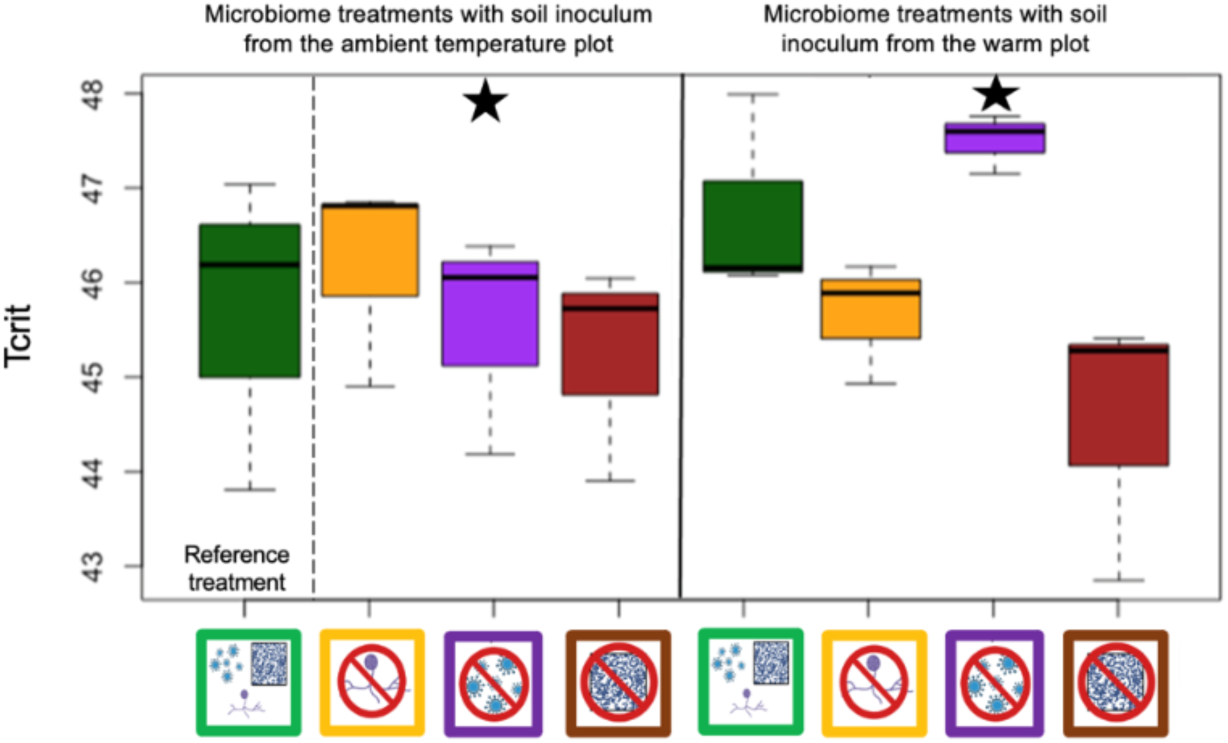
*Tcrit* plotted as a function of the interaction between Treatment and Soil source. Green bar: unaltered microbiomes; yellow bar: reduced AMF; purple bar: reduced fungal pathogens; brown bar: reduced microbes. The star indicates a significant difference between reduced fungal pathogens in the ambient temperature compared to in the warmed plot soil source.

## Discussion

Leaf heat tolerance is a trait that provides valuable insight into the capacity of tropical plants to withstand higher temperatures. Using an *in-situ* tropical forest warming experiment, we showed that high temperatures impacts might become increasingly detrimental if *Guarea guidonia* experiences high leaf temperature. To investigate further the indirect effects of warming through changes in soil microbiome, we used a shade house experiment manipulating soil inoculum from the field experiment. Our treatments successfully altered the soil microbiome in complex ways.

As expected, we found an overall negative effect of fungal pathogens on leaf heat tolerance as seedlings from the reduced fungal pathogens showed a higher *Tcrit* value and *F_V_/F_m_* values over the other treatments. Additionally, the soil microbiome effects on LHT were inconsistent between soil sources. Our results suggest that the soil microbiome might become increasingly essential to buffer plants against climate change, as the soil microbiome significantly influenced *F_V_/F_m_* before *T50* for seedlings growing with soil inoculum from the warmed plot.

### In the field, warming lowers leaf heat tolerance despite a potential for acclimation

In the warmed plots, plants exhibited low leaf heat tolerance despite signs of acclimation at ambient temperature (25°C). Under 25°C, *F_V_/F_m_* was significantly higher in plants in the warmed plots compared to the ambient plots. This result aligns with studies showing that long-term acclimation to elevated temperatures increases the LHT (Baürle, 2016; Friedrich et al., 2019; Filaček et al., 2022). Our results are also consistent with previous physiological studies of acclimation at the TRACE site, where shrubs of *Psychotria brachiata* under *in-situ* experimental warming demonstrated the ability to enhance photosynthesis by broadening its thermal niche and increasing its optimum photosynthetic temperature in response to warmer temperatures (Carter et al., 2020). However, at the same site *Guarea guidonia* did not acclimate to *in-situ* warming (Carter et al., 2018; Carter et al., 2020; Doughty et al., 2023). Yet, these previous studies at TRACE focused on adult trees (Carter et al., 2018; Carter et al., 2020; Doughty et al., 2023) whereas we measured leaf heat tolerance on seedlings. Individual plants experience drastic changes in their environment as they mature, which results in leaves with different characteristics in seedlings compared to adult trees (Lawrence et al., 2022). As a result, tolerance to stress, including heat stress, is likely to change during a plant lifespan (Lawrence-Paul & Lasky, 2024). Physiological differences with ontogeny could explain why, at the same sites, we found evidence of acclimation in seedlings, but others found no sign of acclimation at the adult stage in the same species (Carter et al., 2018; Carter et al., 2020).

The benefit of increasing *F_V_/F_m_* in the warmed plots disappeared when the leaf disks were incubated at high temperatures (46 and 52 °C). At these temperatures, leaves collected from the ambient plots showed higher *F_V_/F_m_* values than those collected from the warmed plots. Several mechanisms could explain the drop in *F_V_/F_m_*. First, plants in the warmed plots might have reached their maximal leaf heat tolerance; therefore, any increase in temperature becomes detrimental. Another possible explanation for this result at high temperatures is that plants in the warmed plots also experienced water stress. Prior to the hurricanes, the warming treatment significantly reduced soil moisture at the TRACE site (Kimball et al., 2018), however, soil moisture is more variable across the treatments post-hurricane (Cavaleri et al., in prep). Low soil moisture could lower LHT because drought conditions undermine plants’ ability to deal with heat by reducing NADP+ concentrations and leading to the breakdown of the oxygen-evolving complex (Ronde et al., 2004; Siddiqui et al., 2019; Perdomo et al., 1996). Consistent with this idea, a previous study at our site found that soil moisture better predicted daily physiological responses to heat than vegetation temperature in two understory shrub species (Carter et al., 2020). Since plants in the warmed plots are experiencing a drier environment due to the warming treatment (Kimball et al., 2018), they might respond poorly to the added stress of high temperature and exhibit a decrease in *F_V_/F_m_*. Therefore, if plants in the warmed plots have a slight advantage under ambient conditions, further increases in air temperature could have dramatic effects on them in part due to ongoing water stress. Given the climatic predictions that Puerto Rico will experience a warmer and drier climate than it currently is (National Weather Puerto Rico Station WXJ69), future studies should investigate how tropical forests will respond simultaneously to these stressors.

### In the shade house, the soil microbiome treatments influenced fungal community composition in complex ways

In the field, DNA analysis of soil fungi has established that warming treatments have impacted fungal composition. However, the results were only marginally significant possibly due to our small sample size to avoid large plot disturbances (only one sample per field plot). While fungal classes known for being the most fungal pathogens, such as *Dothideomycetes, Euromycetes*, and *Sordariomycets* (Ohm et al., 2012; Covo, 2020; Wang et al., 2023) are similarly present in both inoculants. *Glomeromycetes*, an arbuscular mycorrhizal fungal class, were most prevalent in ambient temperature plots. In contrast, *Archaeosporomycetes*, another arbuscular mycorrhizal fungal class, was only present in the warmed plots. While both classes might show a varied distribution along environmental gradients (Stürmer et al., 2018; Pinto-Figueroa et al., 2019), *Archaeosporomycetes* abundance may be influenced by both climatic and edaphic factors (Pinto-Figueroa et al., 2019)

Analyses of soil fungal DNA from the shade house confirmed that the microbiome treatments altered fungal composition. Specifically, in the microbiome treatments with reduced AMF, reduced fungal pathogens, and reduced microbes, the relative abundance of *Eurotiomycetes* and *Sordariomycetes* —classes including many plant pathogens (Morgan & Kamoun, 2007; Zhang et al., 2006)—were diminished. Likewise, the *Dothideomycetes* class, which includes plant pathogens that can significantly affect agricultural crops (Haridas et al., 2020), had a smaller relative abundance in the reduced fungal pathogens and reduced AMF treatment. Given the number of unknown fungi at the class level in our soil (21.28 %), other fungi important for LHT may be present in our soil even though they have not been identified and linked to LHT yet. Finally, the absence of *Glomeromycetes* in the reduced AMF treatment confirms that the treatment successfully removed the presence of these symbionts. However, the reduced AMF treatment also influenced several fungal pathogens, such as *Dothideomycetes, Eurotiomycetes*, and *Sordariomycetes*. This impact on both beneficial and harmful fungi could explain the lack of a significant effect of this treatment on LHT parameters. Future studies of the impacts of AMF on LHT should consider reducing AMF by preventing root colonization manually using mesh rather than chemically to limit off-target effects.

### In the shade house, the soil microbiome significantly influences Tcrit

Consistent with our prediction, reducing fungal pathogens resulted in plants exhibiting a higher heat-leaf tolerance, suggesting that pathogens negatively affect LHT. Specifically, *Tcrit* was significantly higher in plants grown with reduced fungal pathogens in the warm soil source. Reduced pathogen treatment results in seedlings being better able to deal with heat stress. Plants’ vulnerability to diseases may restrict the energy available to sustain a high level of LHT (Matyssek et al., 2005; Smith et al., 2007). Although the mechanisms by which fungal pathogens alter leaf heat tolerance have not been fully elucidated, they may involve intricate interactions between defense responses, signaling pathways, and the ability of plants to adapt to simultaneous or sequential exposure to both heat stress and pathogen attack (Suzuki & Katano, 2018). Detecting the effect of reduced fungal pathogens on leaf heat tolerance only in a warmed soil source implies essential differences between the two soil sources. We detected a significant interaction between soil source and microbiome treatment on the composition of the fungal community. For example, the soil source from the warmed plot inoculum inoculant exhibited a smaller relative abundance of potential pathogenic class *Eurotiomyvetes* (Covo, 2020). This decrease in fungal pathogens in this soil source and microbiome treatment could potentially explain why the plants exhibited a higher LHT. Fewer pathogens may help the plants create a higher heat tolerance due to the plants’ better allocation of resources to deal with heat stress response.

### In the shade house, the influence of soil microbiome on LHT changes under warming conditions

*F_V_/F_m_* differed significantly across the soil microbiomes in seedlings grown in ambient plot soil at high temperatures (50, 52, and at 54 °C). Contrary to our expectations, seedlings from the reduced AMF treatment group showed higher *F_V_/F_m_* than those grown with an unaltered microbiome. Contrary to our expectations, these results suggested that Banrot fungicide treatment may have removed more harmful fungi than beneficial mycorrhizal fungi. Soil DNA data does indeed confirm that the reduced AMF treatment in ambient plot soil source removed AMF alongside pathogen fungi from *Sordariomycetes* and *Dothideomycetes* classes when compared to the unaltered treatment in ambient plot soil source.

*F_V_/F_m_* was also significantly affected by the soil microbiome from the warmed plot soil source but for heat stress at lower temperatures (46 and 48 °C). As expected, seedlings grown with reduced AMF and reduced microbes had lower *F_V_/F_m_* (which means a lower tolerance to heat stress) than seedlings grown with an unaltered soil microbiome. This result aligns with previous studies showing that beneficial microbes and fungi, such as mycorrhizal fungi, might enhance the LHT (Hubbard et al., 2014; Shekhawat et al., 2022). Consistent with our predictions, we found that seedlings grown with reduced fungal pathogens had higher *F_V_/F_m_* than those grown with an unaltered soil microbiome. Therefore, healthier seedlings could better allocate resources to combat heat damage (Chini et al., 2004).

Our study emphasizes the importance of the soil microbiome in buffering plants against climate change. Seedlings from the ambient plot soil source exhibited significant differences in *F_V_/F_m_* across soil microbiome treatments post-*T50* mean values. Conversely, significant differences across soil microbiome treatments manifested as pre-*T50* mean values for seedlings originating from the warmed plot soil source. The *T50* threshold is the temperature at which *F_V_/F_m_* is reduced by 50%. Recently, *T50* has been found an adaptation to extreme leaf temperatures (Perez and Feeley 2020). If the soil microbiome can modulate *F_V_/F_m_* post-T50, this suggests that the soil microbiome may play an important role in the ability of plants respond to climate change and potentially further adapt to changing temperature.

The need to investigate how plant-microbiome linkages change under climate change is increasingly recognized (Cao et al., 2020; Nottingham et al., 2022). Previous studies have highlighted how the soil microbiome can help plants tolerate drought (Yang et al., 2009; Ortíz et al., 2015; Ngumbi et al., 2016). Our study provides additional evidence on how the soil microbiome becomes increasingly important for plants to respond to warmer temperatures. Modeling studies have made global predictions suggesting an overall increase in the relative abundance and diversity of fungal plant pathogens (Delgado-Baquerizo et al., 2020; Li et al., 2023). Our results indicate that pathogens significantly reduced leaf heat tolerance of *G. guidonia*, therefore a future where plant fungal pathogens are more abundant and diverse could be associated with more vulnerable plants to extreme leaf temperatures. However, it is important to note that local predictions of increase in fungal plant pathogens abundance and diversity are mixed (Garcia et al. 2019; Morrison et al. 2019), and Puerto Rico is predicted to experience more frequent and intense drought which could reduce the abundance and diversity of plant fungal pathogens.

### Conclusion

In conclusion, our results highlight the importance of the soil microbiome on leaf heat tolerance in the context of climate change. Contrary to our expectations, *T50*, a commonly used parameter to characterize leaf heat tolerance, remained stable between warmed and ambient plots in the field and across microbiome treatments in the shade house. However, differences emerged in *Tcrit* as well as the raw *F_V_/F_m_* values. Notably, the impacts of soil microbiome on leaf heat tolerance were exacerbated in soil from our warmed plots, suggesting that the soil microbiome will become increasingly important with climate change. Our study strongly suggests the need for additional studies in the role of the microbiome in regulating LHT in the context of climate change. We have focused on the soil microbiomes, but future research should include the leaf microbiomes which directly impacts leaf function.

## Acknowledgments

This research was funded by NSF grant DEB 2120085, Department of Energy (DE-SC0012000, DE-SC-0011806, 89243018S-SC-000014, 89243018S-SC-000017, DE-SC-0018942, DE-SC0022095, 89243021S-SC-000076) and NSF (DEB-1754713). Support was additionally provided by Oklahoma State University, the USDA Forest Service International Institute of Tropical Forestry, and the University of Puerto Rico – Río Piedras. The USDA Forest Service, the US Department of Energy Office of Science, and the National Science Foundation fund TRACE. We thank the TRACE technical team, William Mejia, Deyaneira Iglesias, Laura Rubio, Katherine Pagan-Rivera, and Alberto Ibanez, for helping with the shade house experiment setup and maintenance. We also thank Dr. Bachelot Lab’s members Shelby McMahan, Elizabeth Struble, and Addison Darby for helping in the DNA extraction process. The authors declare no conflicts of interest.

## Author contributions

GHV and BB designed the LHT study. BB, TEW, and JRL designed the shade house experiment. TEW, SCR, and MAC designed and run the warming experiment. TEW, IGP, GHV, PB, JM set up and managed the shade house experiment. TEW and IGP managed students, experiment, and data collection locally. GHV collected and analyzed LHT data. GHV wrote the first draft and following iterations of the manuscripts. Every author provided feedback and comments.

## Supporting Information

**Figure S1:**
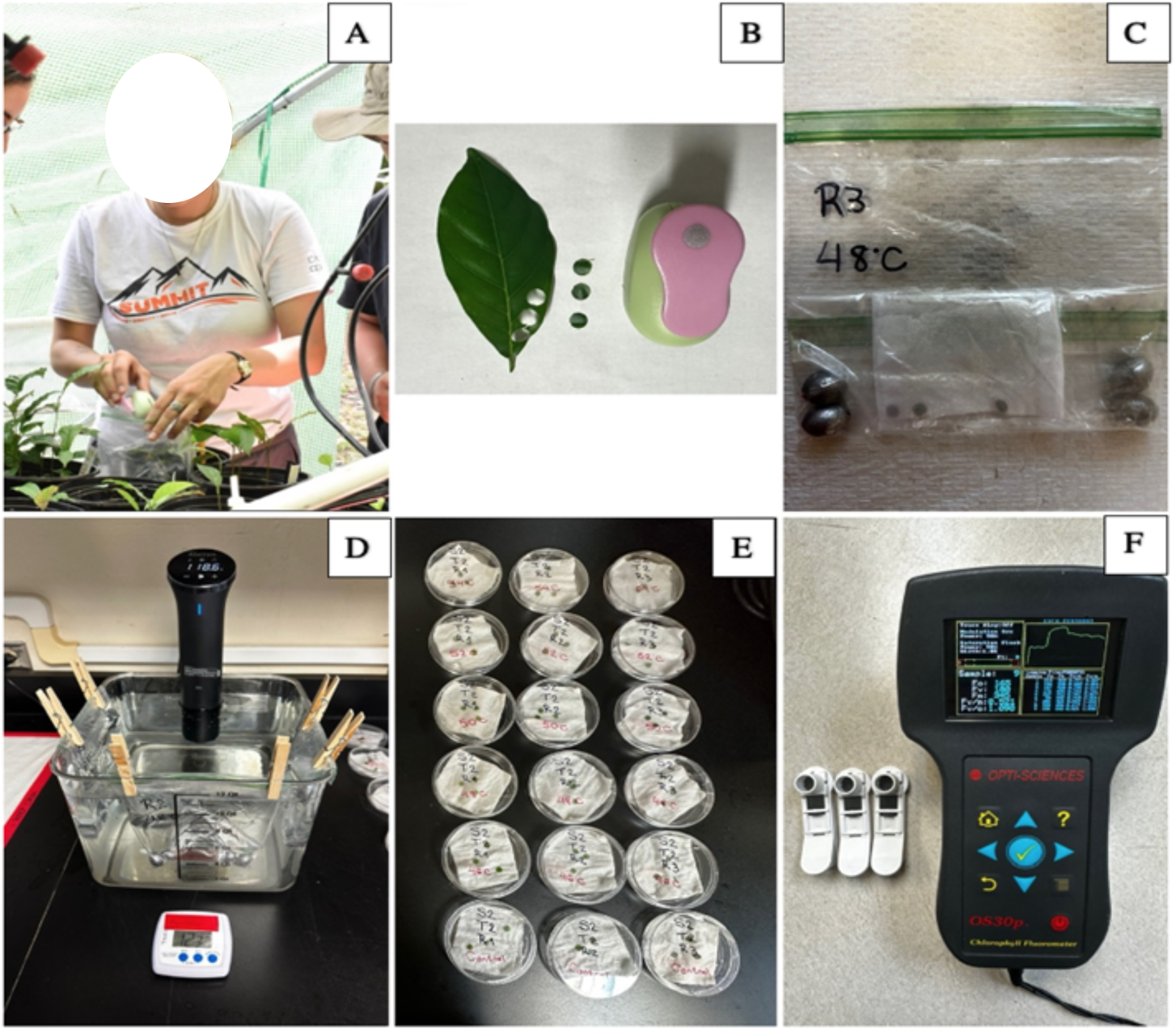
Protocol steps: (A) Collect leaf material (Photo of lead author); (B) Punch leaf disks with a paper puncher; (C) Place leaf disks in a cloth and, then, in Ziploc bags (eggs sinkers used to keep the bag sunk in the water); (D) Apply heat stress in heat controlled water baths for 15 min; (E) Place leaf disks on moisture paper in Petri dishes and wait for 24h; (F) Measure *FV/Fm* with a chlorophyll fluorometer after dark adapt the leaf disks in the clips for about 15 min.

**Figure S2:**
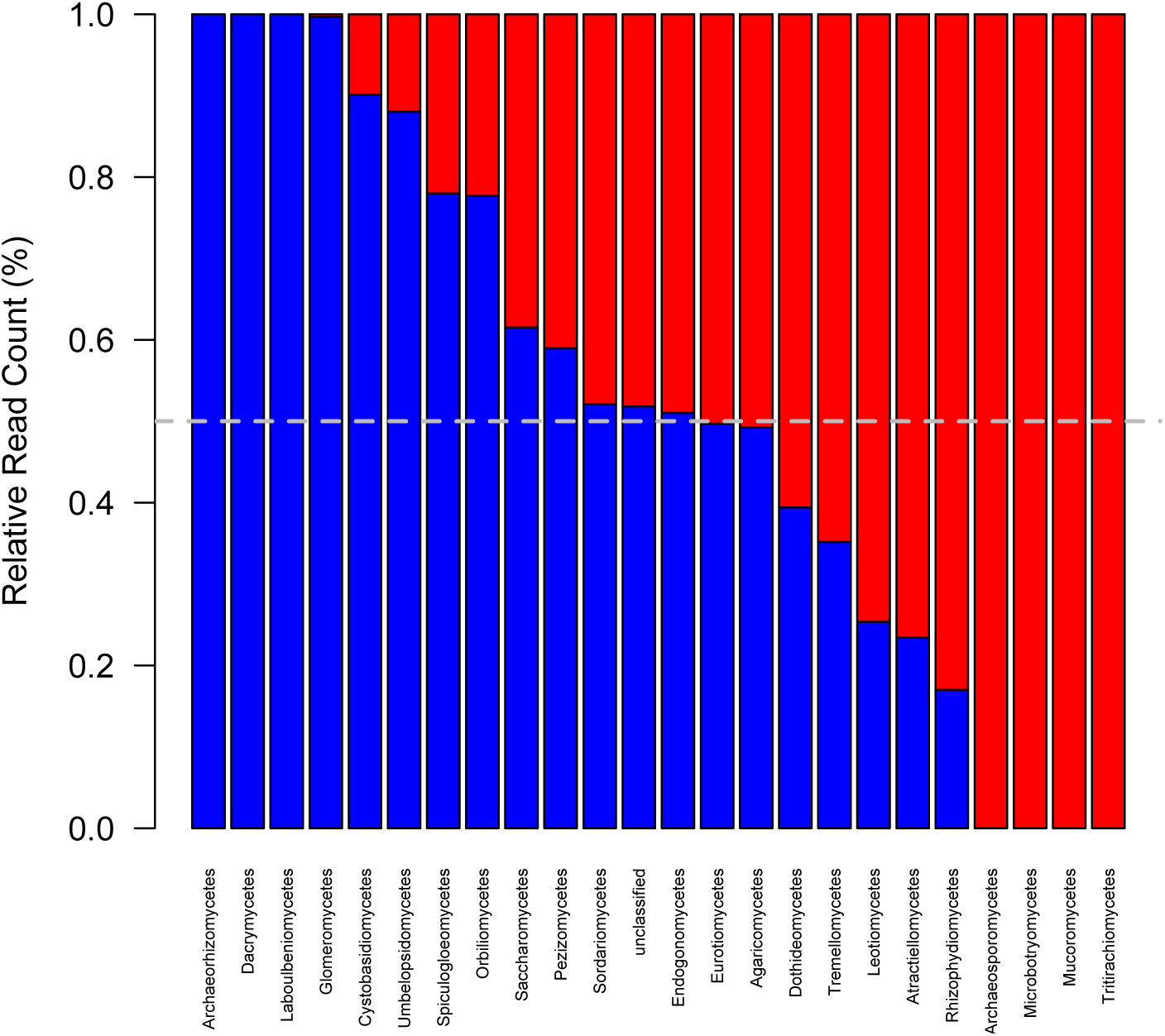
Log Relative abundance (number of reads), standardized to 1, of Fungal Classes in inoculum from the Ambient temperature (blue) or Warmed (red) plots. Dash line is plotted at 50% relative read count.

**Figure S3:**
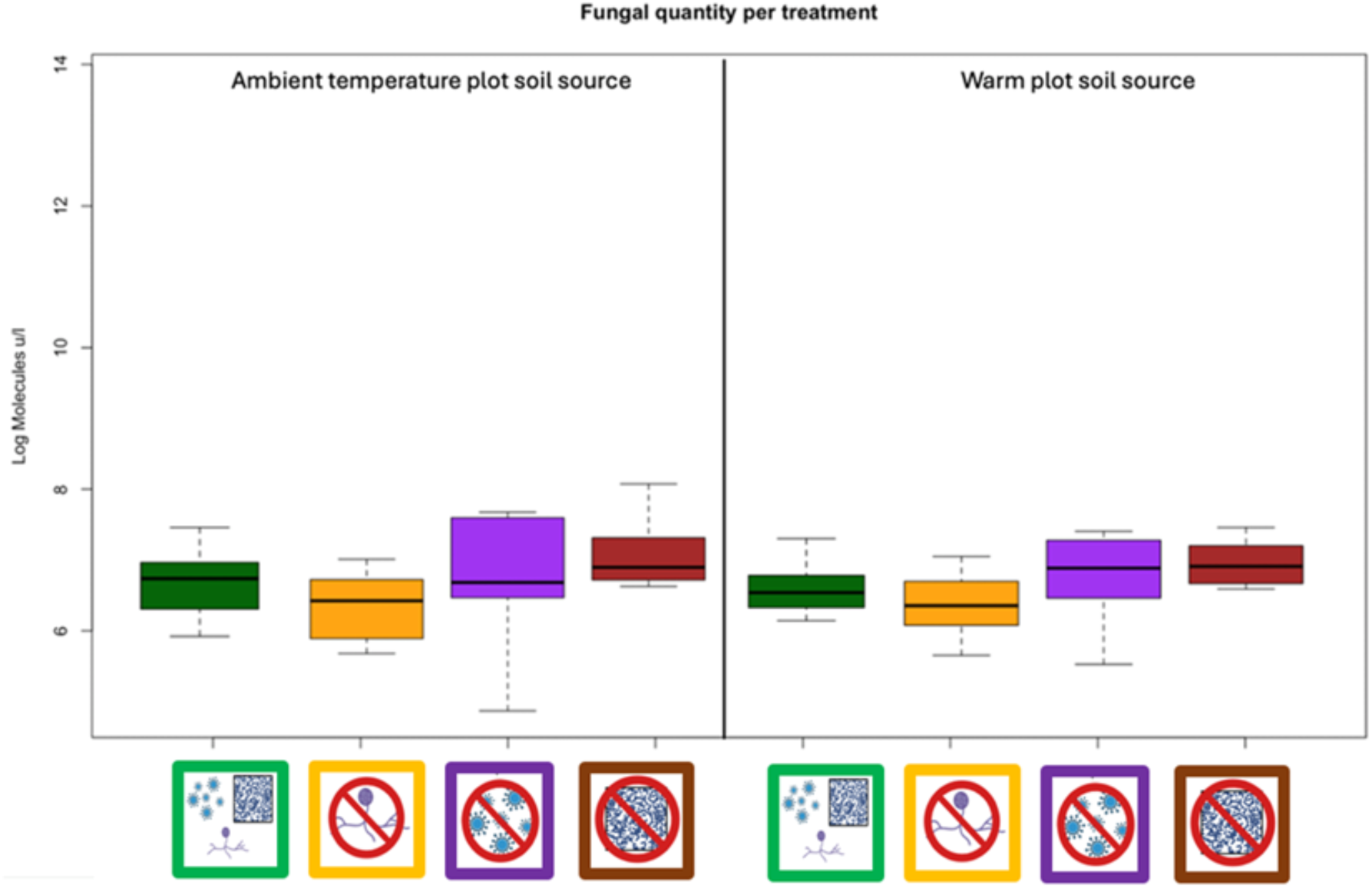
Fungal quantity per treatment in the ambient temperature plot soil source and warmed plot soil source. Green bar: unaltered microbiomes; yellow bar: reduced AMF; purple bar: reduced fungal pathogens; brown bar: reduced microbes.

## REFERENCES

Abarenkov, K., Nilsson, R. H., Larsson, K.-H., Taylor, A. F. S., May, T. W., Frøslev, T. G., Pawlowska, J., Lindahl, B., Põldmaa, K., Truong, C., Vu, D., Hosoya, T., Niskanen, T., Piirmann, T., Ivanov, F., Zirk, A., Peterson, M., Cheeke, T. E., Ishigami, Y.,…Kõljalg, U. (2024). The UNITE database for molecular identification and taxonomic communication of fungi and other eukaryotes: Sequences, taxa and classifications reconsidered. Nucleic Acids Research, 52(D1), D791–D797. 10.1093/nar/gkad1039

Abd El-Daim, I., Bejai, S., & Meijer, J. (2019). Bacillus velezensis 5113 Induced Metabolic and Molecular Reprogramming during Abiotic Stress Tolerance in Wheat. Scientific Reports, 9(1), 1–18. 10.1038/s41598-019-52567-x.

Alonso-Rodríguez, A. M., Wood, T. E., Torres-Díaz, J., Cavaleri, M. A., Reed, S. C., & Bachelot, B. (2022). Understory plant communities show resistance to drought, hurricanes, and experimental warming in a wet tropical forest. Frontiers in Forests and Global Change, 5.

Bale, J. S., Masters, G. J., Hodkinson, I. D., Awmack, C., Bezemer, T. M., Brown, V. K., Butterfield, J., Buse, A., Coulson, J. C., Farrar, J., G. Good, J. E., Harrington, R., Hartley, S., Jones, T. H., Lindroth, R. L., Press, M. C., Symrnioudis, I., Watt, A. D., & Whittaker, J. B. (2002). Herbivory in global climate change research: Direct effects of rising temperature on insect herbivores. Global Change Biology, 8(1), 1-16. 10.1046/j.1365-2486.2002.00451.x

Bachelot, B., Alonso-Rodríguez, A. M., Aldrich-Wolfe, L., Cavaleri, M. A., Reed, S. C., & Wood, T. E. (2020). Altered climate leads to positive density-depend feedbacks in a tropical wet forest. Global Change Biology, 26(6), 3417–3428

Baker, N. R., & Rosenqvist, E. (2004). Applications of chlorophyll fluorescence can improve crop production strategies: an examination of future possibilities. Journal of experimental botany, 55(403), 1607–1621.

Bäurle I. (2016). Plant Heat Adaptation: priming in response to heat stress. F1000Research 5: F1000 Faculty Rev-694.

Bouskill, N. J., Wood, T. E., Baran, R., Zhou, J., Nostrand, J. D. V., Nico, P., Northen, T.R., Silver, W.L. & Brodie, E. L. (2016). Belowground response to drought in a tropical forest soil. I. Changes in microbial functional potential and metabolism. Frontiers in microbiology, 7, 183795.

Carter, K. R., & Cavaleri, M. A. (2018). Within-Canopy Experimental Leaf Warming Induces Photosynthetic Decline Instead of Acclimation in Two Northern Hardwood Species. Frontiers in Forests and Global Change, 1, 423027. 10.3389/ffgc.2018.00011

Carter, K. R., Wood, T. E., Reed, S. C., Schwartz, E. C., Reinsel, M. B., Yang, X., & Cavaleri, M. A. (2020). Photosynthetic and Respiratory Acclimation of Understory Shrubs in Response to in situ Experimental Warming of a Wet Tropical Forest. Frontiers in Forests and Global Change, 3, 576320. 10.3389/ffgc.2020.576320

Cao, J., Teng-Chiu Lin, Zhijie Yang, Yong Zheng, Lin Xie, Decheng Xiong, Yusheng Yang. (2020). Warming exerts a stronger effect than nitrogen addition on the soil arbuscular mycorrhizal fungal community in a young subtropical Cunninghamia lanceolata plantation, Geoderma, Volume 367, 2020,114273, ISSN 0016-7061. 10.1016/j.geoderma.2020.114273.

Chini, A., Grant, J.J., Seki, M., Shinozaki, K. and Loake, G.J. (2004). Drought tolerance established by enhanced expression of the CC–NBS–LRR gene, ADR1, requires salicylic acid, EDS1, and ABI1. The Plant Journal, 38: 810-822. 10.1111/j.1365-313X.2004.02086.x

Cook, B., Corp, L., Nelson, R., Middleton, E., Morton, D., McCorkel, J., Masek, J., Ranson, K., Ly, V., & Montesano, P. (2013). NASA’s LiDAR, hyperspectral and thermal (G-LiHT) airborne imager. Remote Sensing, 5(8), 4045–4066. 10.3390/rs508 4045

Covo, S. (2020). Genomic Instability in Fungal Plant Pathogens. Genes, 11(4), 421. 10.3390/genes11040421

Crous, K. Y., Cheesman, A. W., Middleby, K., Rogers, E. I., Bouet, A. Y., Ellsworth, D. S., Liddell, M. J., Cernusak, L. A., & Barton, C. V. (2023). Similar patterns of leaf temperatures and thermal acclimation to warming in temperate and tropical tree canopies. Tree Physiology, 43(8), 1383–1399. 10.1093/treephys/tpad054

Delgado-Baquerizo, M., Guerra, C. A., Cano-Díaz, C., Egidi, E., Wang, J. T., Eisenhauer, N.,…& Maestre, F. T. (2020). The proportion of soil-borne pathogens increases with warming at the global scale. Nature Climate Change, 10(6), 550–554.

Doughty, C. E., Keany, J. M., Wiebe, B. C., Carter, K. R., Middleby, K. B., Cheesman, A. W., Goulden, M. L., Da Rocha, H. R., Miller, S. D., Malhi, Y., Fauset, S., Gloor, E., Slot, M., Oliveras Menor, I., Crous, K. Y., Goldsmith, G. R., & Fisher, J. B. (2023). Tropical forests are approaching critical temperature thresholds. Nature, 621(7977), 105–111. 10.1038/s41586-023-06391-z

Elad, Y. & Pertot, I. (2014). Climate Change Impacts on Plant Pathogens and Plant Diseases, Journal of Crop Improvement, 28:1, 99–139, DOI: 10.1080/15427528.2014.865412.

Filaček, A., Živčák, M., Ferroni, L., Barboričová, M., Gašparovič, K., Yang, X., Landi, M., & Brestič, M. (2022). Pre-Acclimation to Elevated Temperature Stabilizes the Activity of Photosystem I in Wheat Plants Exposed to an Episode of Severe Heat Stress. Plants, 11(5), 616. 10.3390/plants11050616

Friedrich T, Faivre L, Bäurle I, Schubert D. (2019). Chromatin-based mechanisms of temperature memory in plants. Plant, Cell & Environment 42: 762– 770.

Garcia, M. O., Templer, P. H., Sorensen, P. O., Sanders-DeMott, R., Groffman, P. M., & Bhatnagar, J. M. (2020). Soil microbes trade-off biogeochemical cycling for stress tolerance traits in response to year-round climate change. Frontiers in Microbiology, 11, 616.

Gohl, D. M., Vangay, P., Garbe, J., MacLean, A., Hauge, A., Becker, A., Gould, T. J., Clayton, J. B., Johnson, T. J., Hunter, R., Knights, D., & Beckman, K. B. (2016). Systematic improvement of amplicon marker gene methods for increased accuracy in microbiome studies. Nature Biotechnology, 34(9), 942–949. 10.1038/nbt.3601

Gweon HS, Oliver A, Taylor J, Booth T, Gibbs M, Read DS, Griffiths RI, Schonrogge K. (2015). PIPITS: an automated pipeline for analyses of fungal internal transcribed spacer sequences from the Illumina sequencing platform. Methods in Ecology Evolution, 6(8), 973–980. doi: 10.1111/2041-210X.12399. Epub 2015 May 25. PMID: 27570615; PMCID: PMC4981123.

Haridas, S., Albert, R., Albert, R., Binder, M., Bloem, J., LaButti, K., Salamov, A., Andreopoulos, B., Baker, S., Barry, K., Bills, G., Bluhm, B., Cannon, C., Castanera, R., Castanera, R., Culley, D., Daum, C., Ezra, D., Gonzalez, J., Henrissat, B., Henrissat, B., Henrissat, B., Kuo, A., Liang, C., Lipzen, A., Lutzoni, F., Magnuson, J., Mondo, S., Mondo, S., Nolan, M., Ohm, R., Ohm, R., Pangilinan, J., Park, H., Ramírez, L., Alfaro, M., Sun, H., Tritt, A., Yoshinaga, Y., Zwiers, L., Turgeon, B., Goodwin, S., Spatafora, J., Crous, P., Grigoriev, I., & Grigoriev, I. (2020). 101 Dothideomycetes genomes: A test case for predicting lifestyles and emergence of pathogens. Studies in Mycology, 96, 141–153. 10.1016/j.simyco.2020.01.003.

Heartsill-Scalley, T., Scatena, F. N., Estrada, C., McDowell, W. H., & Lugo, A. E. (2007). Disturbance and long-term patterns of rainfall and throughfall nutrient fluxes in a subtropical wet forest in Puerto Rico. Journal of Hydrology, 333(2-4), 472–485.

Holdridge, L. R. (1967). Life zone ecology. Tropical Science Center, San Jose, Costa Rica.

Hubbard, M., Germida, J.J. and Vujanovic, V. (2014) Fungal endophytes enhance wheat heat and drought tolerance in terms of grain yield and second-generation seed viability. J Appl Microbiol 116, 109–122.

Jiang, Y., & Huang, B. (2001). Effects of calcium on antioxidant activities and water relations associated with heat tolerance in two cool-season grasses. Journal of experimental botany, 52 355, 341–9. 10.1093/JEXBOT/52.355.341.

Kimball, B. A., Alonso-Rodríguez, A. M., Cavaleri, M. A., Reed, S. C., González, G., & Wood, T. E. (2018). Infrared heater system for warming tropical forest understory plants and soils. Ecology and Evolution, 8(4), 1932–1944.

Klockmann, M., Günter, F., & Fischer, K. (2017). Heat resistance throughout ontogeny: body size constrains thermal tolerance. Global Change Biology, 23(2), 686–696.

Krause, G. H., Winter, K., Krause, B., Jahns, P., García, M., Aranda, J., & Virgo, A. (2010). High-temperature tolerance of a tropical tree, Ficus insipida: methodological reassessment and climate change considerations. Functional Plant Biology, 37(9), 890–900.

Krause, G. H., Cheesman, A. W., Winter, K., Krause, B., & Virgo, A. (2013). Thermal tolerance, net CO2 exchange and growth of a tropical tree species, Ficus insipida, cultivated at elevated daytime and nighttime temperatures. Journal of Plant Physiology, 170(9), 822–827.

Kullberg AT, Coombs L, Soria Ahuanari RD, Fortier RP, Feeley KJ. (2024). Leaf thermal safety margins decline at hotter temperatures in a natural warming experiment in the Amazon. New Phytol. 2024 Feb;241(4):1447-1463. doi: 10.1111/nph.19413. Epub 2023 Nov 20. PMID: 37984063.

Kuznetsova, A., Brockhoff, P. B., & Christensen, R. H. B. (2017). lmerTest package: tests in linear mixed effects models. Journal of statistical software, 82(13).

Lawrence, E. H., C. J. Springer, B. R. Helliker, and R. S. Poethig. (2022). The carbon economics of vegetative phase change. Plant Cell and Environment 45: 1286–1297.

Lawrence-Paul, E. H., & Lasky, J. R. (2024). Ontogenetic changes in ecophysiology are an understudied yet important component of plant adaptation. American Journal of Botany, e16294. 10.1002/ajb2.16294

Lenth, R. (2022). emmeans: Estimated marginal means, aka least-squares means. R package version 1.7. 2.

Le Quéré, C., Andrew, R. M., Friedlingstein, P., Sitch, S., Hauck, J., Pongratz, J.,…& Keeling, R. F. (2018). Global carbon budget 2018. Earth System Science Data, 10(1), 405–448.

Li, P., Tedersoo, L., Crowther, T. W., Wang, B., Shi, Y., Kuang, L.,…& Jiang, J. (2023). Global diversity and biogeography of potential phytopathogenic fungi in a changing world. Nature Communications, 14(1), 6482.

Matsui, T., & Eguchi, H. (1971). Effects of Environmental Factors on Leaf Temperature In a Temperature Controlled Room. Environment control in biology, 19, 69–72. 10.2525/ECB1963.19.69.

Matyssek, R., Agerer, R., Ernst, D., Munch, J., Osswald, W., Pretzsch, H., Priesack, E., Schnyder, H., & Treutter, D. (2005). Plants’ capacity in regulating resource demand. Plant biology, 7 6, 560–80. 10.1055/S-2005-872981.

Mau A., Reed S.C., Wood T.E., & Cavaleri M.A. (2018). Temperate and tropical forest canopies are already functioning beyond their thermal thresholds for photosynthesis. Forests 9: 47.

Miller B.D., Carter K.R., Reed S.C., Wood T.E., & Cavaleri M.A. (2021). Only sun-lit leaves of the uppermost canopy exceed both air temperature and photosynthetic optima in a wet tropical forest. Agricultural and Forest Meteorology 301–302: 108347.

Morgan, W., & Kamoun, S. (2007). RXLR effectors of plant pathogenic oomycetes. Current opinion in microbiology, 10 4, 332–8. 10.1016/J.MIB.2007.04.005.

Nakmee, P.S.; Techapinyawat, S.; Ngamprasit, S. (2016). Comparative potentials of native arbuscular mycorrhizal fungi to improve nutrient uptake and biomass of Sorghum bicolor Linn. Agric. Nat. Resour. 50, 173–178.

Morrison, E. W., Pringle, A., van Diepen, L. T., Grandy, A. S., Melillo, J. M., & Frey, S. D. (2019). Warming alters fungal communities and litter chemistry with implications for soil carbon stocks. Soil Biology and Biochemistry, 132, 120–130.

National Weather Puerto Rico Station WXJ69. (2024). https://www.weather.gov/nwr/sites?site=WXJ69.

Nelson, E. B. (1994). The disease triangle and the disease cycle. Turf Glass Trends ISSN, 1076-7207.

Ngumbi, E., & Kloepper, J. (2016). Bacterial-mediated drought tolerance: Current and future prospects. Applied Soil Ecology, 105, 109–125. 10.1016/J.APSOIL.2016.04.009.

Nguyen, N. H., Song, Z., Bates, S. T., Branco, S., Tedersoo, L., Menke, J., Schilling, J. S., & Kennedy, P. G. (2016). FUNGuild: An open annotation tool for parsing fungal community datasets by ecological guild. Fungal Ecology, 20, 241–248. 10.1016/j.funeco.2015.06.006

Nottingham, A. T., Whitaker, J., Ostle, N. J., Bardgett, R. D., McNamara, N. P., Fierer, N., Meir, P. (2019). Microbial responses to warming enhance soil carbon loss following translocation across a tropical forest elevation gradient. Ecology Letters, 22(11), 1889–1899.

Nottingham, A. T., Scott, J. J., Saltonstall, K., Broders, K., Montero-Sanchez, M., Püspök, J., Meir, P. (2022). Microbial diversity declines in warmed tropical soil and respiration rise exceed predictions as communities adapt. Nature Microbiology, 7(10), 1650–1660.

Ohm RA, Feau N, Henrissat B, Schoch CL, Horwitz BA, et al. (2012) Diverse Lifestyles and Strategies of Plant Pathogenesis Encoded in the Genomes of Eighteen Dothideomycetes Fungi. PLOS Pathogens 8(12): e1003037. 10.1371/journal.ppat.1003037

Oksanen, J., et al. (2022) Vegan: Community Ecology Package. https://cran.r-project.org/package=vegan.

Ortíz, N., Armada, E., Duque, E., Roldán, A., & Azcón, R. (2015). Contribution of arbuscular mycorrhizal fungi and/or bacteria to enhancing plant drought tolerance under natural soil conditions: effectiveness of autochthonous or allochthonous strains. Journal of plant physiology, 174, 87–96. 10.1016/j.jplph.2014.08.019.

Parry, M.L., Canziani, O.F., Palutikof, J.P., van der Linden PJ, Hanson CE (eds). (2007). Climate change 2007: Impacts, Adaptation and Vulnerability. Contribution of Working Group II to the Fourth Assessment Report of the Intergovernmental Panel on Climate Change Cambridge University Press, Cambridge, UK, 976 pp.

Padfield, D., & Matheson, G. (2018). nls.multstart: Robust Non-Linear Regression using AIC Scores. R package version 1.0.0. Retrieved from https://CRAN.R-project.org/package=nls.multstart.

Pan, Y., Birdsey, R.A., Phillips, O.L., Jackson, R.B. (2013). The structure, distribution, and biomass of the world’s forests. Ann Rev Ecol Evol System 44:593–622. 10.1146/annurev-ecolsys-110512-135914.

Perdomo, P., Murphy, J. A., & Berkowitz, G. A. (1996). Physiological changes associated with performance of Kentucky bluegrass cultivars during summer stress. HortScience, 31(7), 1182–1186.

Perez, T. M., & Feeley, K. J. (2020). Photosynthetic heat tolerances and extreme leaf temperatures. Functional Ecology, 34(11), 2236–2245.

Pennington, T. D., & Clarkson, J. J. (2013). A revision of Guarea (Meliaceae). Edinburgh Journal of Botany, 70(2), 179–362.

Perera, R., Cullen, B., & Eckard, R. (2019). Using Leaf Temperature to Improve Simulation of Heat and Drought Stresses in a Biophysical Model. Plants, 9. 10.3390/plants9010008.

Pinto-Figueroa, E., Seddon, E., Yashiro, E., Buri, A., Niculita-Hirzel, H., Meer, J., & Guisan, A. (2019). Archaeorhizomycetes Spatial Distribution in Soils Along Wide Elevational and Environmental Gradients Reveal Co-abundance Patterns With Other Fungal Saprobes and Potential Weathering Capacities. Frontiers in Microbiology, 10. 10.3389/fmicb.2019.00656.

Prabhakaran, A. D., & Dann, E. K. (2022). Evaluation of fungicide soil drench treatments to manage black root rot disease of avocado. Plant Disease, 106(8), 2026–2030.

Ramakrishna Akula & Gokare Aswathanarayana Ravishankar. (2011). Influence of abiotic stress signals on secondary metabolites in plants, Plant Signaling & Behavior, 6:11, 1720–1731, DOI: 10.4161/psb.6.11.17613

Reed, S. C., Reibold, R., Cavaleri, M. A., Alonso-Rodríguez, A. M., Berberich, M. E., & Wood, T. E. (2020). Soil biogeochemical responses of a tropical forest to warming and hurricane disturbance. Advances in Ecological Research, 62, 225–252. 10.1016/bs.aecr.2020.01.007

R Core Team. (2022). R: A language and environment for statistical computing. Vienna, Austria: R Foundation for Statistical Computing. Retrieved from https://www.R-project.org.

Ronde, J., Cress, W., Krüger, G., Strasser, R., & Staden, J. (2004). Photosynthetic response of transgenic soybean plants, containing an Arabidopsis P5CR gene, during heat and drought stress. Journal of plant physiology, 161 11, 1211–24. 10.1016/J.JPLPH.2004.01.014.

Sardans, J., Hans Lambers, C. Preece, A. Fahad A., & Penuelas J. (2023). Role of mycorrhizas and root exudates in plant uptake of soil nutrients (calcium, iron, magnesium, and potassium): has the puzzle been completely solved? The Plant Journal, 114 (6), 1227–1242.

Sauer, D. B., & Burroughs R. (1986) Disinfection of seed surfaces with sodium hypochlorite. Phytopathology 76 (7), 745–749.

Scatena, F. N. (1989). An introduction to the physiography and history of the Bisley Experimental Watersheds in the Luquillo Mountains of Puerto Rico (Vol. 72). US Department of Agriculture, Forest Service, Southern Forest Experiment Station.

Sentinella, A. T., Warton, D. I., Sherwin, W. B., Offord, C. A., & Moles, A. T. (2020). Tropical plants do not have narrower temperature tolerances but are more at risk from warming because they are close to their upper thermal limits. Global Ecology and Biogeography, 29(8), 1387–1398. 10.1111/geb.13117

Shekhawat, K., Almeida-Trapp, M., García-Ramírez, G. X., & Hirt, H. (2022). Beat the heat: Plant-and microbe-mediated strategies for crop thermotolerance. Trends in Plant Science.

Slot, M., Krause, G. H., Krause, B., Hernández, G. G., & Winter, K. (2019). Photosynthetic heat tolerance of shade and sun leaves of three tropical tree species. Photosynthesis Research, 141, 119–130.

Smith, V. (2007). Host resource supplies influence the dynamics and outcomes of infectious diseases. Integrative and comparative biology, 47 2, 310–6. 10.1093/icb/icm006.

Siddiqui, Z.A.; Pichtel, J. (2008). Mycorrhizae: An Overview. In Mycorrhizae: Sustainable Agriculture and Forestry; Springer Science and Business Media LLC: Berlin, Germany, 2008; pp. 1–35.

Solly, E., Lindahl, B., Dawes, M., Peter, M., Souza, R., Rixen, C., & Hagedorn, F. (2017). Experimental soil warming shifts the fungal community composition at the alpine treeline. The New Phytologist, 215 2, 766–778. 10.1111/nph.14603.

Schloss PD et al. (2009). Introducing mothur: Open-source, platform-independent, community-supported software for describing and comparing microbial communities. Applied and Environmental Microbiology 75:7537–7541.

Starkey, T. E., Enebak, S. A., McQuage, K., & Barfield, K. (2013). Control of Rhizoctonia foliar blight in forest seedling nurseries: a 3-year study. National Proceedings: Forest and Conservation Nursery Associations–2012, 11.

Suzuki, N.; Katano, K. (2018). Coordination Between ROS Regulatory Systems and Other Pathways Under Heat Stress and Pathogen Attack. Front. Plant Sci. 2018, 9.

Tedersoo, L.; Sánchez-Ramírez, S.; Kõljalg, U.; Bahram, M.; Döring, M.; Schigel, D.S.; May, T.; Ryberg, M.; Abarenkov, K. (2018). High-level classification of the Fungi and a tool for evolutionary ecological analyses. Fungal Divers. 2018, 90, 135–159.

Stürmer, S. L., Bever, J. D., & Morton, J. B. (2018). Biogeography of arbuscular mycorrhizal fungi (Glomeromycota): a phylogenetic perspective on species distribution patterns. Mycorrhiza, 28(7), 587–603.

Tarvainen, L., Wittemann, M., Mujawamariya, M., Manishimwe, A., Zibera, E., Ntirugulirwa, B.,…& Uddling, J. (2022). Handling the heat–photosynthetic thermal stress in tropical trees. New Phytologist, 233(1), 236–250.

Teskey, R., Wertin, T., Bauweraerts, I., Ameye, M., Mcguire, M.A., Steppe, K. (2015). Responses of tree species to heat waves and extreme heat events. Plant, Cell & Environment, 38 (2015), pp. 1699–1712. 10.1111/pce.12417.

Tiwari, R., Gloor, E., da Cruz, W. J. A., Schwantes Marimon, B., Marimon-Junior, B. H., Reis, S. M.,…& Galbraith, D. (2021). Photosynthetic quantum efficiency in south-eastern Amazonian trees may be already affected by climate change. Plant, Cell & Environment, 44(7), 2428–2439.

Tunison R.P., Wood T.E., Reed S.C., Cavaleri M.A. (2023) Respiratory acclimation of tropical forest roots in response to in situ experimental warming and hurricane disturbance. Ecosystems. 10.1007/s10021-023-00880-y

Velásquez, A.C., Castroverde, C.D.M., He, S.Y. (2018). Plant-Pathogen Warfare under Changing Climate Conditions. Curr Biol. 2018 May 21;28(10): R619-R634. doi: 10.1016/j.cub.2018.03.054. PMID: 29787730; PMCID: PMC5967643.

Wang, Z., Kim, W., Wang, Y.-W., Yakubovich, E., Dong, C., Trail, F., Townsend, J. P., & Oded Yarden. (2023). The Sordariomycetes: an expanding resource with Big Data for mining in evolutionary genomics and transcriptomics. 4. 10.3389/ffunb.2023.1214537

Waqas, M., Khan, A. L., Shahzad, R., Ullah, I., Khan, A. R., & Lee, J. (2015). Mutualistic fungal endophytes produce phytohormones and organic acids that promote japonica rice plant growth under prolonged heat stress. Journal of Zhejiang University. Science. B, 16(12), 1011–1018. 10.1631/jzus.B1500081

Wilson, G. W. T., & Williamson, M. M. (2008). Topsin-M: the new benomyl for mycorrhizal-suppression experiments. Mycologia, 100(4), 548–554.

Wood, T. E., Cavaleri, M. A., Giardina, C. P., Khan, S., Mohan, J. E., Nottingham, A. T., Reed, S. C., & Slot, M. (2019). Soil warming effects on tropical forests with highly weathered soils. Ecosystem Consequences of Soil Warming, 385-439. 10.1016/B978-0-12-813493-1.00015-6

Yaffar, D., Wood, T. E., Reed, S. C., Branoff, B. L., Cavaleri, M. A., & Norby, R. J. (2021). Experimental warming and its legacy effects on root dynamics following two hurricane disturbances in a wet tropical forest. Global Change Biology, 27(24), 6423–6435. 10.1111/gcb.15870

Yang, J., Kloepper, J., & Ryu, C. (2009). Rhizosphere bacteria help plants tolerate abiotic stress. Trends in plant science, 14 1, 1–4. 10.1016/j.tplants.2008.10.004.

Zhang, N., Castlebury, L., Miller, A., Huhndorf, S., Schoch, C., Seifert, K., Rossman, A., Rogers, J., Kohlmeyer, J., Volkmann-Kohlmeyer, B., & Sung, G. (2006). An overview of the systematics of the Sordariomycetes based on a four-gene phylogeny. Mycologia, 98 6, 1076–87. 10.3852/MYCOLOGIA.98.6.1076.

